# Beyond adoption rates: Farmer motivations and communication needs in straw management decision-making

**DOI:** 10.1101/2025.08.25.672071

**Authors:** Kristina Blennow, Elin Anander

## Abstract

Sustainable bioenergy is central to climate change mitigation, yet biomass supply depends not only on biophysical and economic assessments but also on farmers’ decision-making. Straw from cereal and oilseed crops can support renewable energy, but its availability is constrained by on-farm uses, management practices, and farmers’ access to knowledge of sustainable soil management. Because straw removal affects soil organic carbon (SOC) stocks, which underpins soil fertility, nutrient cycling, and carbon sequestration, understanding farmers’ motivations and informational needs is critical to support sustainable straw management.

We examined the determinants of straw management among farmers in Scania County, southern Sweden, testing three hypotheses: that manure application increases the likelihood of straw removal, that a higher proportion of leased land promotes straw removal, and that humus-rich soils are associated with a higher probability of straw removal. Survey data from 2021 were analysed using Bayesian Additive Regression Trees (BART) and permutation tests, incorporating farmers’ expected utility.

Results show that manure application increases removal probability, whereas a higher share of leased land reduces it. Removal was most frequent on soils with intermediate humus content. Cluster analysis identified three farmer profiles, revealing heterogeneous motivations and knowledge gaps, particularly regarding soil humus content, and underscoring the need for context-specific communication strategies.

These findings demonstrate that similar straw management behaviours can arise from diverse motivations and local conditions. Linking expectations, soil properties, and practices provides actionable insights for targeted advisory and policy measures that balance agronomic and economic outcomes with long-term SOC-mediated fertility, carbon sequestration, and ecosystem services.

## 1. Introduction

According to the Intergovernmental Panel on Climate Change (IPCC), sustainable bioenergy is among the low-carbon energy sources projected to expand significantly in most mitigation pathways that limit global warming to 1.5–2°C, contributing to the rapid decarbonisation of the energy supply sector (Riahi et al. 2022). In alignment with this global mitigation context, the European Union (EU) has established targets through the European Green Deal (European Council 2025) and the revised Renewable Energy Directive (RED III), which specify that at least 42.5% of the EU’s gross final energy consumption should come from renewable sources by 2030 (EU 2023a). Biomass, currently the largest source of renewable energy in the EU (EEA 2025), is anticipated to contribute to these targets by replacing fossil fuels in energy production and serving as a feedstock for bio-based materials and products.

The potential to expand biomass production sustainably depends on aligning technical, environmental, and economic assessments with practical decision-making by landowners, particularly farmers. Straw from cereal and oilseed crops is identified in policy frameworks as a potential feedstock due to its high productivity and limited direct competition with food production (Imperial College 2021). Despite technical assessments indicating substantial biophysical potentials, the market availability of straw remains limited. Previous research suggests that straw is commonly used within farming systems, for example, as animal bedding, which reduces its availability for external markets (Townsend et al. 2018; Blennow et al. 2025a).

The limited incorporation of farmers’ perspectives in biomass potential assessments can result in inaccurate estimates of available biomass supply and policy expectations. Studies have documented that market uncertainties and price sensitivity can influence farmers’ willingness to remove or sell biomass (Giannoccaro et al. 2017; Gérard and Jayet 2023; Thomson Ek 2024), although this relationship is not consistent across contexts. For instance, Blennow et al. (2025a) reported that the supply of cereal and oilseed straw was more strongly associated with factors such as land tenure and straw end-use than with price. Land tenure reflects farmers’ incentives and constraints related to soil fertility management. Farmers who lease a large share of their land may operate with shorter planning horizons or have weaker incentives to invest in long-term soil fertility. Since manure use is an important factor in maintaining soil organic matter, it can also shape decision-making on whether to retain or remove straw. Understanding how these factors interact with soil fertility management is therefore critical for interpreting straw management practices and their implications for sustainable biomass supply.

Straw management is not only a matter of nutrient supply and farming systems but is also central to maintaining soil organic carbon (SOC), which sustains soil fertility through improved structure, water retention, and nutrient cycling. Both straw use for bioenergy and straw retention for SOC are motivated by climate change mitigation: while bioenergy substitutes fossil fuels, initiatives such as the 4 per mille program (Minasny et al. 2017) stress that increasing SOC stocks by 0.4% annually could offset global CO₂ emissions over 20 years (Martin et al. 2021). Farmers’ choices on straw removal or retention thus influence both soil fertility and climate mitigation, yet bioenergy strategies rarely address their informational needs or decision-making contexts (e.g., Scarlat et al. 2019).

This study identifies the motivations and communication needs of farmers in southern Sweden regarding the potential supply of cereal and oilseed straw for energy and bio-based materials. Using a problem-feeding approach (Thorén and Persson 2013; Persson et al. 2018), challenges in practical agriculture were identified and investigated through a survey (Blennow et al. 2025a). By analysing decision-making, we assess factors associated with straw removal and the types of information and policy support farmers need.

Declining SOC levels in Swedish agricultural soils raise concerns about sustainability (Barrios Latorre et al. 2024a). Simulations show that safe removal rates vary spatially (Barrios Latorre et al. 2024b). Farmers’ expected impacts on fertility strongly influence decision-making; for example, 90% of Dutch farmers intend to increase soil organic matter (SOM) (Hijbeek et al. 2018). Although SOM and SOC are not identical, they are closely correlated, suggesting similar patterns in Sweden: farmers managing soils with higher humus content may be more confident in removing straw, whereas those with lower SOC may retain the straw to protect fertility. Conversely, farmers with lower SOC levels may retain straw to protect soil fertility.

Specifically, we test the following hypotheses:

1. The likelihood of straw harvest removal is higher among farmers who apply manure compared to those who do not (*cf.* Blennow et al. 2025a).
2. A higher proportion of leased land is positively associated with straw harvest removal decision-making (*cf.* Blennow et al. 2025a).
3. Farmers managing humus-rich soils are more likely to engage in straw harvest removal decision-making than farmers managing soils with lower humus content (*cf.* Hijbeek et al. 2018).

The decision-making analysis applied in this study builds on a framework developed by Persson et al. (2020) and Blennow et al. (2020), further described in Blennow et al. (2021). This framework, which has previously been applied to studies of decision-making in climate change mitigation and adaptation (Blennow and Persson 2021; Blennow et al. 2025b) and the cultivation of *Populus* spp. (Blennow et al. 2025c) uses survey data to assess respondents’ expected utility (*sensu* von Neumann and Morgenstern 1944) when deciding on the supply of biomass from cereal and oilseed straw.

## 2. Materials and methods

### 2.1 Study area

Scania County, located in southern Sweden (Figure 1), is distinguished by a substantial proportion of agricultural land, encompassing approximately 45% of its total area (Statistics Sweden 2023), and by a high concentration of agricultural enterprises. The county’s arable soils are among the most fertile in Sweden, supporting intensive production of annual crops, primarily cereals (Swedish Board of Agriculture 2025). Notably, land use patterns exhibit considerable heterogeneity across the region: in the lowland coastal areas, agricultural land constitutes 60–81% of the landscape, whereas the northern central areas are predominantly forested, with agricultural land accounting for only 5–19%. The intermediate zone between these areas is characterised by a more balanced distribution of forest and farmland (Swedish Board of Agriculture 2025) (Figure 1).

**Figure 1.**
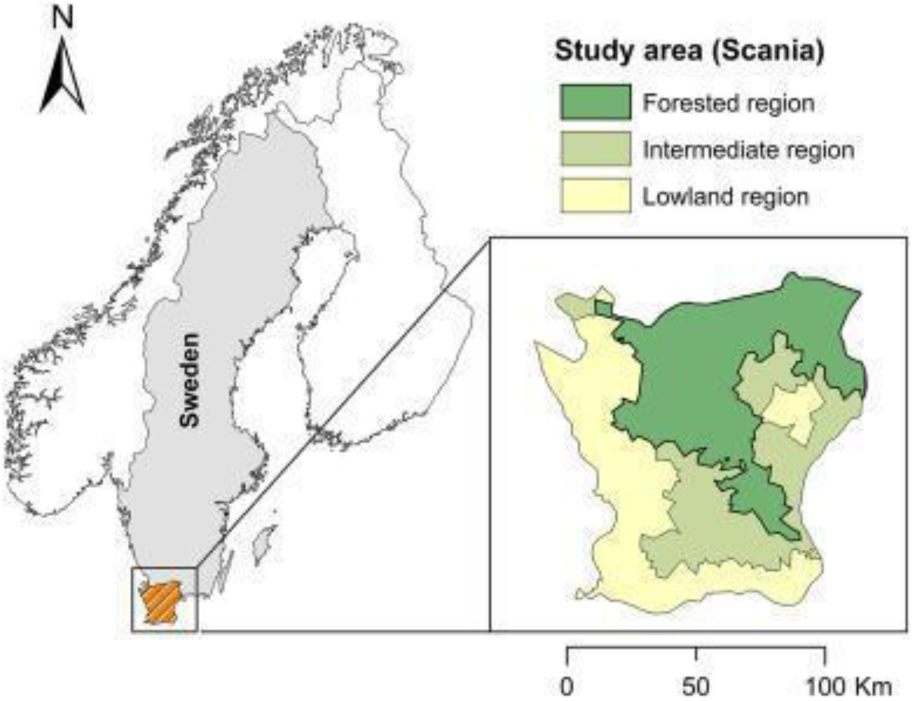
The study area, Scania, is located in southern Sweden. The colour coding indicates three dominant types of land use. Made with Natural Earth and reproduced from Anander et al. (2024).

Animal husbandry, particularly cattle and dairy production, is predominantly concentrated in the forested and intermediate zones, while it is relatively uncommon in the lowland areas (Erlandsson 1999). In 2020, the total agricultural area in Scania encompassed 490,000 hectares, of which arable land, excluding pasture, accounted for 435,000 hectares, corresponding to 17% of Sweden’s total arable area of 2.5 million hectares (Swedish Board of Agriculture 2025). Furthermore, Scania represents the largest contributor to Sweden’s straw biomass potential, providing approximately 40% of the national potential for straw-based energy production (Börjesson 2021).

### 2.2 Data collection

#### 2.2.1 Survey design and sample selection

Building on insights gained from an exploratory pre-study, a broad, unstratified internet-based survey was implemented to capture a diverse sample of farmers (Blennow et al., 2025a). Approximately 50% of members of the Federation of Swedish Farmers (LRF) in Scania were randomly selected and invited via e-mail by LRF to participate. The survey focused on the potential supply of cereal and oilseed straw for energy and material applications, with a total sample size of 2,558 respondents. The parent LRF membership group represents approximately 85% of all agricultural holdings recorded in Scania’s farm register as of 2020 (Swedish Board of Agriculture 2025).

The survey was administered in Scania using the Netigate platform (2024) for questionnaire development and distribution. Participants received a cover letter detailing the study’s objectives and clarifying the voluntary nature of participation (see Blennow et al. 2025a).

#### 2.2.2. Data collection and response rate

The survey was launched on February 17, 2021 and remained open until April 23, 2021. One reminder was sent to all farmers in the sample to encourage participation.

The questionnaire consisted of 35 questions and is described in detail in Blennow et al. (2025a). For this study, responses to ten questions were analysed (Table 1). These questions pertain to farm location and size, soil humus content, manure application, removal of cereal and oilseed straw, and respondents’ expected impacts of cereal and oilseed straw removal.

**Table 1.** Questions analysed in the present study on the removal of cereal and oilseed straw. See Blennow et al. (2025a) for the complete questionnaire.

| Number | Question | Response option |
| --- | --- | --- |
| Q.1 | In which harvest zone is the land you cultivate located?<sup>a</sup> | Aggregated into three regions:<br>Forested region<br>Intermediate region<br>Lowland region<br>I do not know |
| Q.2 | How many ha of arable land does the farm you manage encompass? (Include both owned and leased arable land, but exclude arable land you lease out.) | It does not have any arable land<br>It has _____ ha of arable land |
| Q.3 | How many ha of the arable land is leased? | I do not lease any arable land<br>I lease _____ ha of arable land |
| Q.4 | What humus content would you say is most common in the arable land you cultivate? | a. Less than 2%<br>b. 2 - 3%<br>c. 3 - 4%<br>d. 4 - 5%<br>e. 5 - 6%<br>f. More than 6%<br>g. Don't know the humus content |
| Q.5 | On the farm you cultivated in 2020: Did you remove any straw from cereals or oil crops that was used for energy production (e.g., for heat or biogas production, either used on your farm or sold externally), on or off-farm livestock use (e.g. bedding for animals) or other off-farm use?<sup>b</sup> | a/ Yes<br>b/ No<br>c/ I did not grow cereals or oil crops in 2020 |
| Q.6 | Did you apply any of the following to the arable land you cultivated during 2020?<br>(Tick the option(s) that apply for the year 2020)<sup>c</sup> | a/ Manure from own animals<br>b/ Manure from external livestock<br>c/ Sludge |
| Q.7 | On your farm, do you expect that straw removal<br>• worsens the farm's economic profitability?<br>• worsens soil structure?<br>• reduces soil humus content?<br>• negatively affects the establishment of subsequent crops?<br>• disrupts workflow on the farm?<br>• makes the soil harder to till?<br>• increases weed problems? | Yes, always<br>Often<br>Seldom<br>No, never<br>Don't know |
| Q.8 | On your farm, do you expect that straw removal<br>• improves the farm's economic profitability?<br>• improves soil structure?<br>• improves soil humus content?<br>• positively affects the establishment of subsequent crops? | Yes, always<br>Often<br>Seldom<br>No, never<br>Don't know |
|  | • facilitates workflow on the farm?<br>• makes the soil easier to till?<br>• reduces weed problems? |  |

<sup>a</sup> The detailed farm locations were initially collected based on multiple harvest zones. For this analysis, these zones were aggregated into three broader regions: Forested region, Intermediate region, and Lowland region (see Anander et al. 2024).
<sup>b</sup> Constructed by aggregating three survey questions.
<sup>c</sup> Additional response options in the original question, specifically the two sludge alternatives (sewage sludge from wastewater treatment plants and digestate from biogas production), were combined into a single category, while other options were excluded from the analysis because they did not contain humus.

A total of 174 responses were received, corresponding to a response rate of 7%. Of these, 110 respondents had completed Q.7 and Q.8 in Table 1, corresponding to a response rate of 4%. After excluding 16 respondents who had not cultivated cereals and/or oilseed crops in 2020, 94 questionnaires remained for analysis, representing approximately 2% of the total farming community in Scania.

### 2.3. Data preparation

In line with the methodology described by Persson et al. (2020) and Blennow et al. (2020), we calculated the Net Value of Expected Impacts (NVEI) for the decision to remove cereal and oilseed straw from arable land (see Q.5 in Table 1). The NVEI was derived by summing the net scores of paired positive and negative expectations associated with each biomass provision decision (see Q.7 and Q.8 in Table 1).

For each expectation, responses of “Yes, always” were assigned a value of +4 (for positive expectations) or −4 (for negative expectations), while “Often” responses were assigned +3 or −3, respectively. All other response options (e.g., “Seldom,” “Never,” “Don’t know”) were scored as 0. These scores were aggregated across relevant items for each respondent to calculate the overall NVEI score. Partial NVEI scores, ranging from -4 to 4, were also computed to quantify the contribution of individual sub-questions (i.e., specific expectations).

### 2.4 Statistical analysis and machine learning modelling

Bayesian methods were chosen for their robustness in handling small and unevenly distributed datasets. Unlike frequentist techniques, Bayesian inference provides credible parameter estimates and posterior probabilities even with limited sample sizes (McElreath 2016). This approach also enables direct probabilistic statements about group differences. All analyses were conducted in R version 4.4.1 (R Core Team, 2024).

To assess the strength of evidence, we applied Jeffreys’ (1939) interpretation scale (see Table 2). Weakly informative priors were incorporated to stabilise the estimation process, ensuring minimal impact on posterior distributions and preserving clarity following Jeffreys’ scale.

**Table 2.** Strength of evidence based on posterior probability (Jeffreys 1939).

| Evidence strength | Posterior probability |
| --- | --- |
| Null hypothesis supported | $< 0.5$ |
| Bare mention | $0.50 \leq p < 0.77$ |
| Substantial evidence | $0.77 \leq p < 0.91$ |
| Strong evidence | $0.91 \leq p < 0.97$ |
| Very strong evidence | $0.97 \leq p < 0.99$ |
| Decisive evidence | $\geq 0.99$ |

#### 2.4.1 Machine learning modelling

We applied Bayesian Additive Regression Trees (BART) to model the probability of removing cereal and/or oilseed straw in 2020 (*Removal*, Q.5 in Table 1), using NVEI scores as a predictor, with the *bartMachine* package in R (Kapelner and Bleich 2016). BART is a non-parametric machine learning technique that constructs a collection of shallow regression trees, with each tree adding incrementally to the overall prediction. The method uses a Bayesian approach to regularise the model and reduce the risk of overfitting. To further assess model performance and guard against overfitting, we applied 5-fold cross-validation, in which the data were randomly split into five subsets, with each subset used once as a test set while the remaining four subsets were used for training. The probabilistic outputs of BART were interpreted as the probability of choosing to remove straw.

Additional predictors were included to test the hypotheses and to guide the identification of communication needs: the share of leased land (*Lease share*), manure application (*Manure,* including manure from own animals and external livestock), and the predominant humus content of farm soils (*Humus content*).

Model performance was evaluated using the Area Under the Curve (AUC) of the Receiver Operating Characteristic (ROC), representing the probability that a model ranks a randomly chosen positive instance higher than a randomly chosen negative instance. Model interpretation was supported by Partial Dependence Plots (PDPs) generated using the *ICEbox* package (Goldstein et al., 2015).

#### 2.4.2 Hypothesis testing and effect direction

To assess the statistical significance of individual predictors in the BART model, we performed permutation tests using the *bartMachine* package in R (Kapelner et al. 2016). This method involves repeatedly shuffling the values of a predictor variable while keeping all other variables fixed and recalculating the model’s performance for each permutation. The observed difference in model performance is then compared to the distribution of differences obtained under the null hypothesis (random permutations), allowing computation of a p-value. Predictors with p-values less than the significance threshold of α = 0.05 were considered statistically significant. This approach provides a robust, non-parametric way to test predictor importance without relying on distributional assumptions.

However, permutation tests do not indicate whether effects are positive or negative. Therefore, to determine the direction and magnitude of the relationships, we calculated SHAP (SHapley Additive exPlanations) values using the *iml* package (Molnar et al. 2018), visualised with the *shapviz* package (Mayer 2025). SHAP values decompose each prediction into additive contributions from predictors, indicating whether a predictor increases or decreases the probability of straw removal for each observation.

To examine differences between groups of respondents, we conducted Bayesian proportion tests. This test was performed using a non-informative prior with a prior probability of 0.5 and was implemented using the R package *Bayesian First Aid* (Bååth 2004).

#### 2.4.3 Identification of communication needs

To identify distinct groups of respondents, we performed cluster analysis directly on the raw dataset. The dichotomous variable *Manure* was included as a binary factor (0 = no, 1 = yes). *Humus content* was treated as a factor, with levels 5 and 6 aggregated due to only one observation in level 5 (see Q.4 in Table 1). *Lease share* ranged from 0 to 1.

A Gower distance matrix was calculated to accommodate the mixed variable types. We explored different clustering approaches, including Partitioning Around Medoids (PAM) and hierarchical clustering with Ward’s method, and evaluated the optimal number of clusters using average silhouette widths using the *cluster* package (Maechler et al. 2025). A choice of three clusters was made based on interpretability and silhouette scores.

The hierarchical clustering method was applied to the Gower distance matrix using the *hclust* package (base R), and cluster assignments were extracted for the chosen number of clusters. Cluster cohesion and separation were visualised using silhouette plots using the *factoextra* package (Kasambara and Mundt 2020).

Cluster profiles were generated by calculating the mean values of all variables within each cluster, except for *Humus content*, for which the modal value was used. Heatmaps were used to visualise cluster characteristics using the package *ggplot2* (Wickham 2016): main variables such as *Farm profitability, Soil structure, Humus content effect, Subsequent crops, Work flow, Tilling properties,* and *Weed* were shown in one heatmap, while *Manure*, *Lease share,* and *Humus content* were visualised separately to highlight relative scale differences.

Associations between removal status (0/1) and partial NVEI variables were analysed using Bayesian Gaussian regression models in *brms* (Bürkner 2017). For each cluster, each NVEI variable was modelled as a Gaussian outcome predicted by removal status. Posterior distributions were then used to estimate the expected differences between respondents who removed and those who did not, providing a probabilistic measure of these differences that incorporates uncertainty in the estimates.

## 3 Results

### 3.1 Modelling cereal and oilseed straw removal

#### 3.2.1. Univariate modelling with NVEI

Sixty out of 94 respondents reported having decided to remove cereal and oilseed straw in 2020 (Q.5 in Table 1). To explore the usefulness of the NVEI as a predictor, a univariate BART model was fitted. The univariate model demonstrated a reasonable ability to classify farmers’ decisions, achieving an Area Under the Curve (AUC) of 0.77 (Figure S1). This result indicates that NVEI captures relevant aspects of farmers’ decision-making processes. The model revealed an almost steadily increasing probability of deciding to remove straw with increasing NVEI, plateauing at higher positive values (Figure 2).

**Figure 2.**
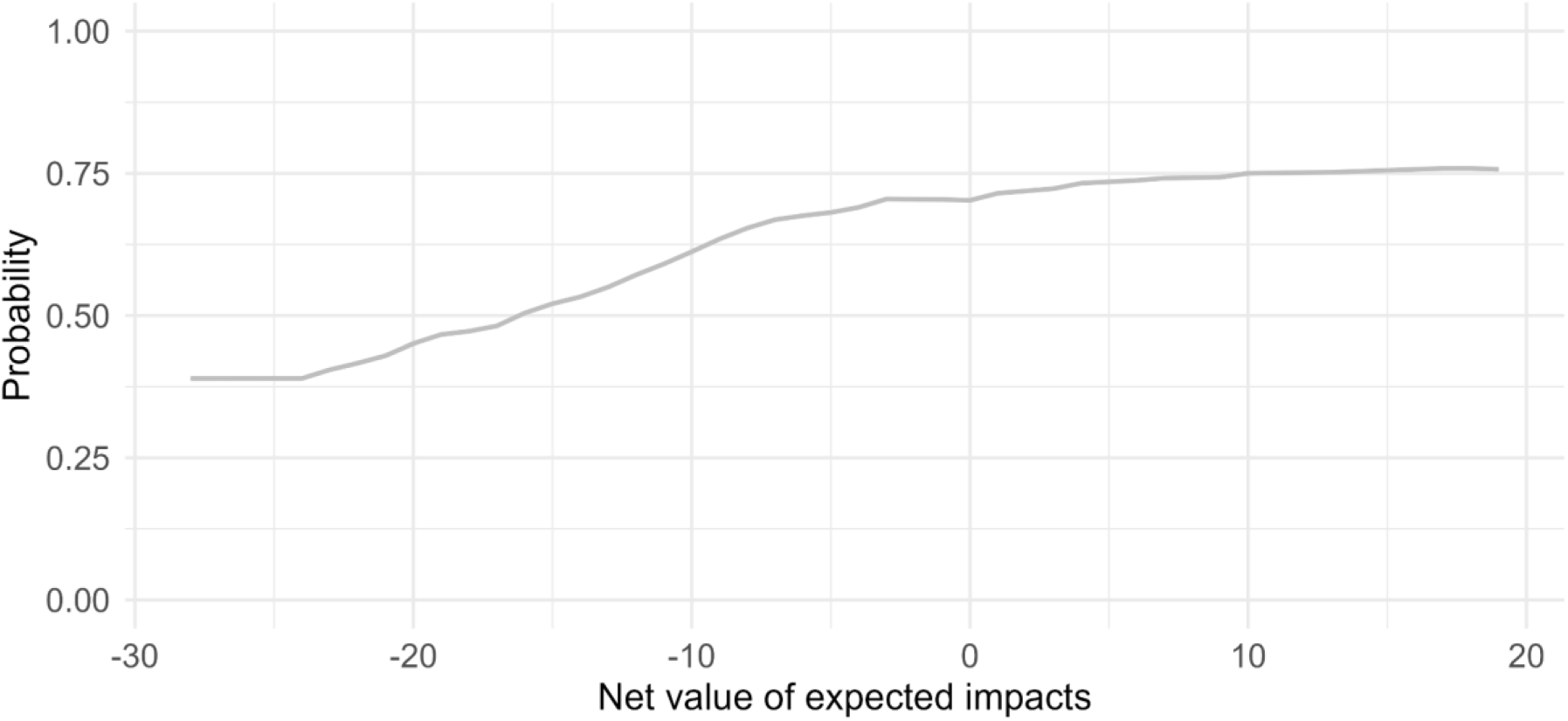
Partial Dependence Curve (PDP) of the relationship between the Net Value of Expected Impact (NVEI) and the predicted probability of having decided to remove cereal and oilseed straw in 2020 based on a BART model (Q.5, 7 and 8 in Table 1).

#### 3.1.2 Multivariate modelling with additional predictors

Building on the insights from the univariate model, a multivariate BART model was fitted, incorporating predictors such as the share of leased land (*Lease share*), the practice of manure application on agricultural land (*Manure*), predominant humus content of farm soils (*Humus content*), and *NVEI*. Convergence diagnostics indicated that the BART model converged satisfactorily (Figure S2). The multivariate model demonstrated an excellent ability to separate the classes, achieving an AUC of 0.90 (Figure S1). The model correctly classified 93% of the positive cases (sensitivity) and 71% of the negative cases (specificity) (Table 3).

**Table 3.** Confusion matrix for the multivariate model. Rows represent actual classes, and columns represent predicted classes. Model errors are row-wise error rates, and use errors are column-wise error rates.

|  | Predicted 1 | Predicted 0 | Model errors |
| --- | --- | --- | --- |
| Actual 1 | 56 | 4 | 0.067 |
| Actual 0 | 10 | 24 | 0.294 |
| Use errors | 0.152 | 0.143 | 0.149 |

Overall error rates were balanced across predicted classes (use error = 0.149), confirming that the model provided both high discriminatory power (AUC = 0.90) and strong predictive accuracy. A permutation test found that the predictor variable *NVEI* is significantly associated with the decision to remove (Figure 3).

**Figure 3.**
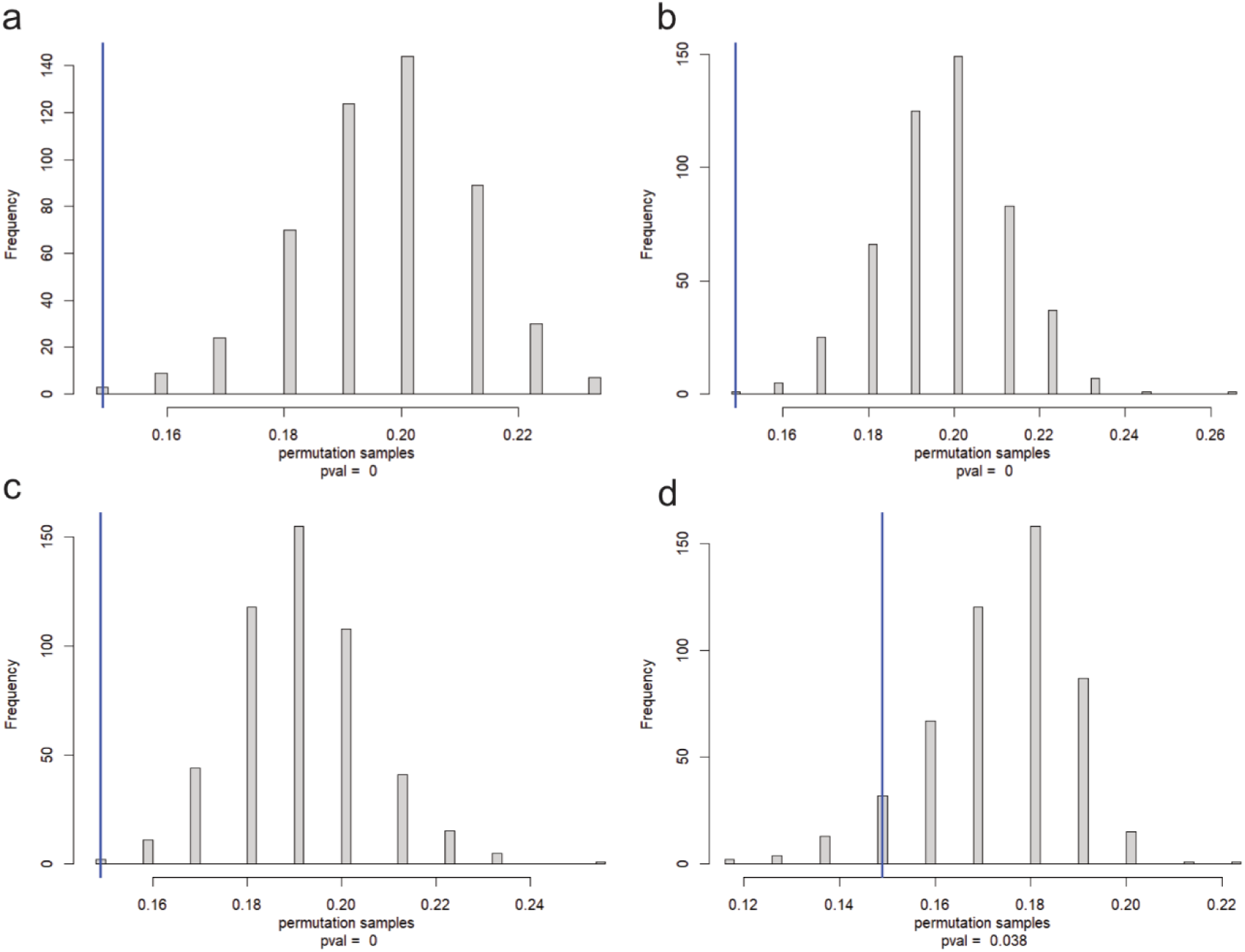
Permutation tests for individual predictor variables used in the multivariate BART model: (a) *NVEI*, (b) *Manure*, (c) *Lease share*, and (d) *Humus content*. Given 500 permutations in each test, a p-value of 0 corresponds to p < 0.002.

### 3.2 Hypothesis testing

#### 3.2.1 H1. The likelihood of straw harvest removal decision-making is higher among farmers who apply manure compared to those who do not

To test H1, which posits that the likelihood of straw harvest removal decision-making is higher among farmers who apply manure compared to those who do not, a permutation test was conducted using a multivariate BART model. The test found that the predictor *Manure* is significantly associated with the decision to remove (p < 0.002; Figure 3). To interpret the influence of individual predictors, SHAP (SHapley Additive exPlanations) values were computed. SHAP values showed that respondents applying manure are more likely to remove straw than those who do not (Figure 4).

**Figure 4.**
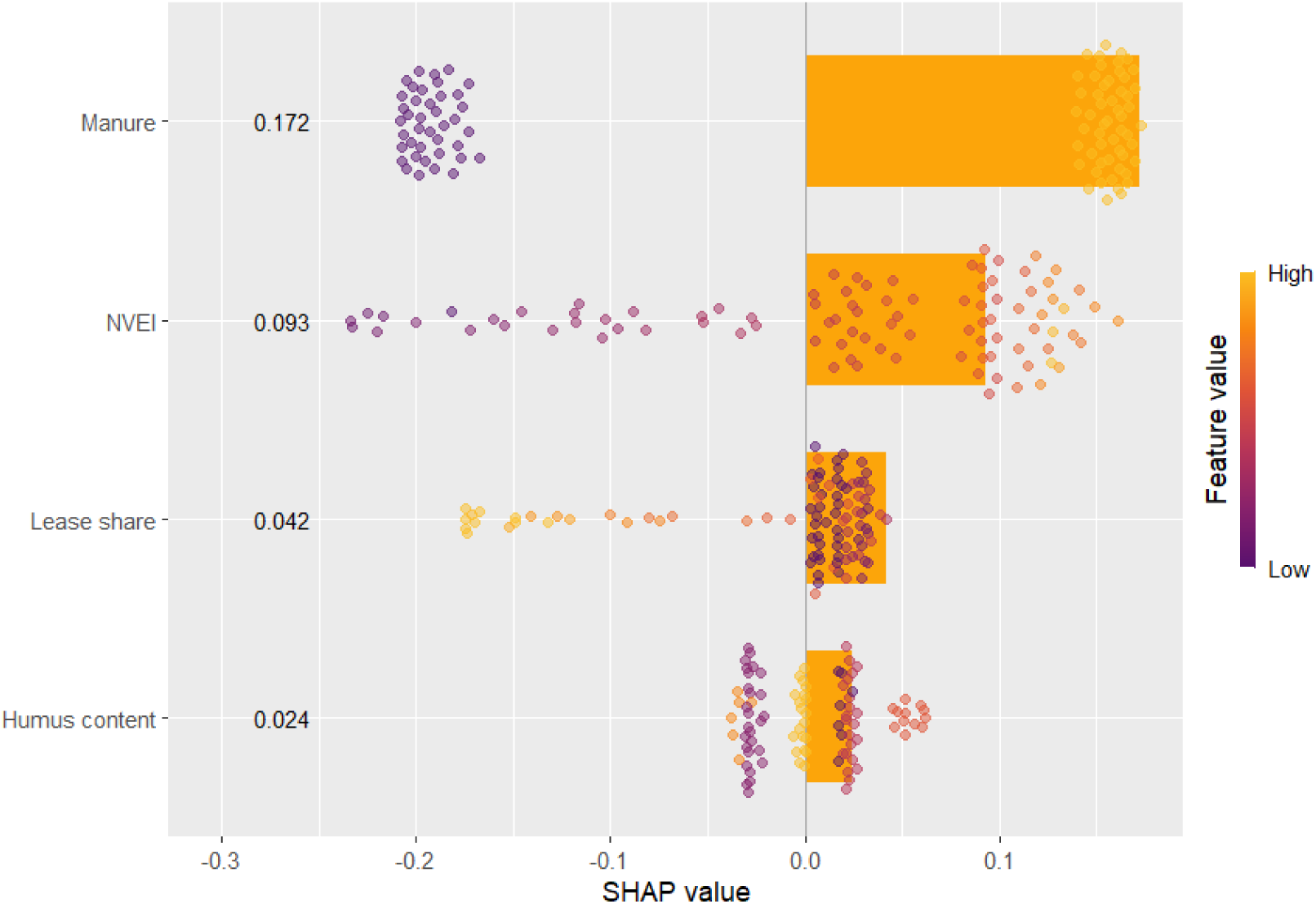
SHAP summary plot showing the relative influence of predictors (manure application, NVEI, share of leased land, and humus content of farm soils) on the probability of deciding to remove cereal and oilseed straw.

#### 3.2.2 H.2 A higher proportion of leased land is associated with straw harvest removal decision-making

To test H.2, which posits that a higher proportion of leased land associated with straw harvest removal decision-making, we employed two complementary approaches. First, we performed an overall Bayesian proportion test that does not account for potential confounding factors.

This comparison provided strong evidence that respondents who did not remove straw in 2020 were more likely to lease more than 50% of their cultivated land than those who did remove it (Table 3). Second, a permutation test based on the multivariate BART model was conducted. The test indicated that the predictor *Lease share* is significantly associated with the decision to remove (p =0.002; Figure 3). A higher share of leased land is associated with a lower probability of cereal and oilseed straw removal (Figure 4).

**Table 3.** Proportion of farmers who did remove straw in 2020 by the share of total arable land area leased, tested using a Bayesian proportion test.

| *Did remove straw in 2020* | *Estimated relative frequency among respondents leasing 50%* | *Comparison (posterior probability)<sup>a</sup>* |
| --- | --- | --- |
| Yes | 0.52 | 6.4% (More than half leased > half or less leased) |
| No | 0.68 | 93.6% (More than half leased > half or less leased) |

<sup>a</sup> Posterior probability reflects the likelihood that respondents who did remove straw in 2020 and leased $\geq 50\%$ of their land exceeds that of respondents leasing $\leq 50\%$

#### 3.2.3 H.3 Farmers managing humus-rich soils are more likely to engage in straw harvest removal decision-making than farmers managing soils with lower humus content

To test H.3, which posits that farmers managing humus-rich soils are more likely to engage in straw harvest removal decision-making than farmers managing soils with lower humus content, a permutation test was conducted using a multivariate BART model. The test found that the predictor *Humus content* is significantly associated with the decision to remove straw (p = 0.04; Figure 3). SHAP values indicated that straw removal was most likely among respondents with intermediate soil humus levels, compared to those with the lowest or highest levels, as well as respondents who were unsure of their soil’s humus content (Figure 4).

### 3.3. Communication needs of respondents regarding removal of cereal and oilseed straw

The multivariate BART model highly accurately predicts removal decision-making among respondents (Table 3). Negative values of NVEI indicate that the respondent perceives the removal of cereal and oilseed straw primarily as a risk, while positive values suggest it is seen as an opportunity (Q.7 and Q.8 in Table 1). The probability of deciding on removal increases with higher NVEI values, reaching a plateau when NVEI becomes positive (Figure 2).

Three main respondent profiles (clusters) were identified based on raw data used in the BART model, with the NVEI variable represented by its partial components (Figure 5).

**Figure 5.**
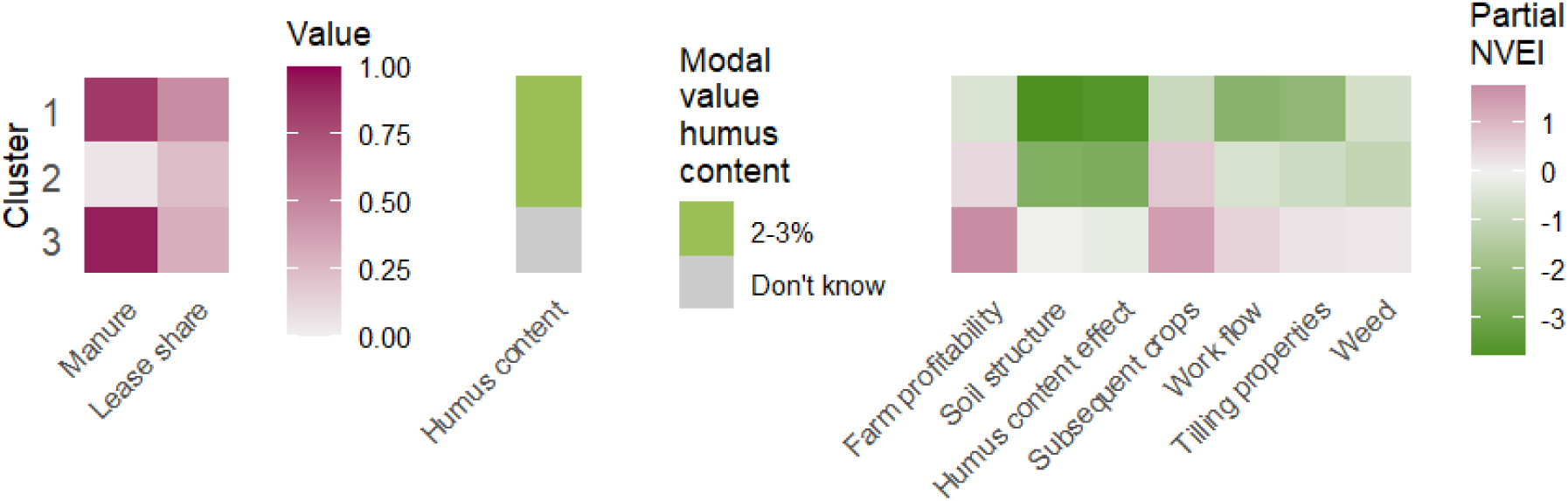
Heatmap of respondent clusters (profiles) based on multiple variables using Gower distance. *Manure* was treated as a binary factor (0 = no, 1 = yes), and *Lease share* ranged from 0 to 1. Mean values are shown for all variables except *Humus content*, which is represented by its modal value. For the NVEI variable, relevant subcomponents were included. All variables used for clustering are displayed in the heatmap.

Using Bayesian regression, posterior probability analysis identified which partial NVEI components were most strongly associated with removal decision-making within each cluster (Figure 6).

**Figure 6.**
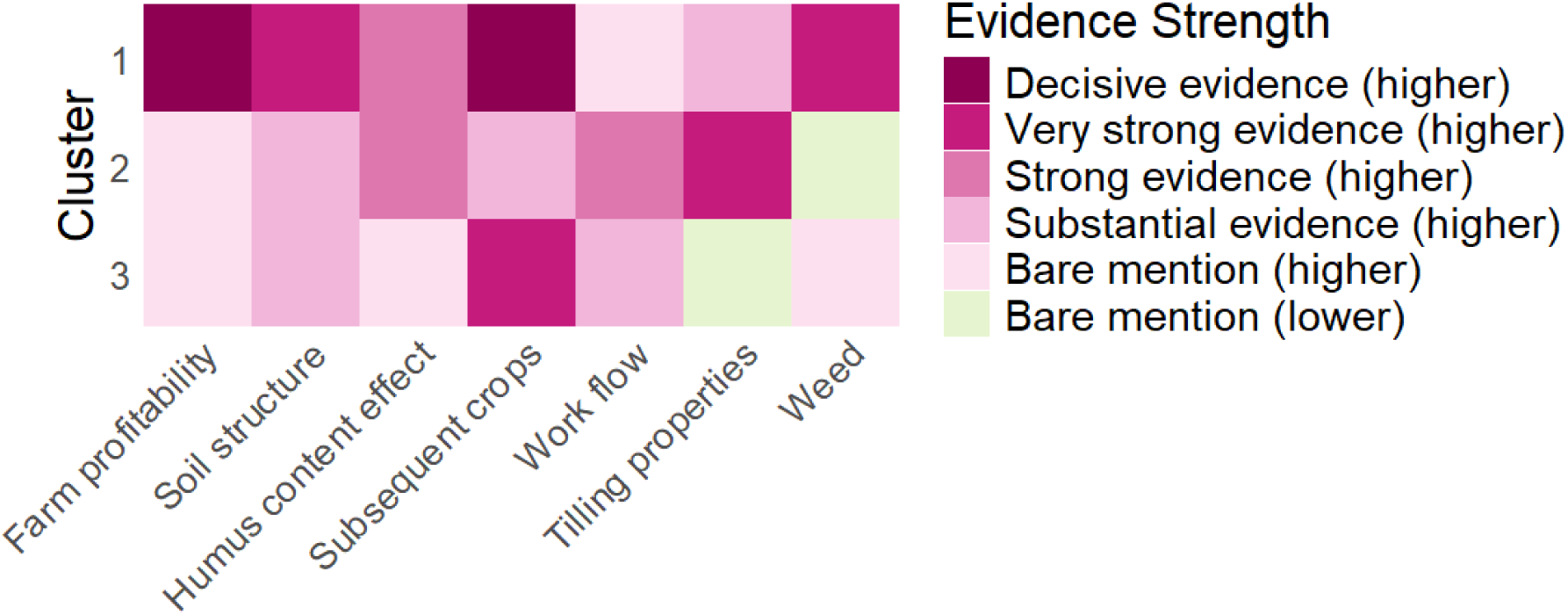
Posterior probabilities of differences in partial NVEI between respondents who did remove straw and those who did not by cluster.

We next examined how the clusters are distributed regionally and how straw removal rates vary between regions. Respondents from the Lowland region were predominantly classified into Cluster 2, whereas none from the Forested region belonged to this cluster (Table 4).

**Table 4.** Number of respondents by region and cluster membership.

| Cluster | Forested<br>(n) | Intermediate<br>(n) | Lowland<br>(n) | Total<br>(per cluster) |
| --- | --- | --- | --- | --- |
| 1 | 2 | 5 | 19 | 26 |
| 2 | 0 | 9 | 26 | 35<sup>a</sup> |
| 3 | 4 | 14 | 13 | 31<sup>a</sup> |

<sup>a</sup> Information on one regional membership is missing.

Differences in straw removal rates across regions were apparent, with generally higher rates observed in Cluster 3 (Table 5). Pairwise comparisons (Figure 7) indicate that in Cluster 1, there is substantial evidence of higher removal rates in the Forested region compared to the Intermediate region, but strong evidence that rates are higher in the Lowland than in the Intermediate region. In Cluster 2, very strong and substantial evidence shows that removal rates are highest in the Lowland region, lower in the Intermediate region, and lowest in the Forested region where no respondent was located. In Cluster 3, there is decisive evidence that removal rates are higher in the Forested region than in the Lowland region, and decisive evidence that removal rates are higher in the Intermediate region than in the Lowland region.

**Figure 7.**
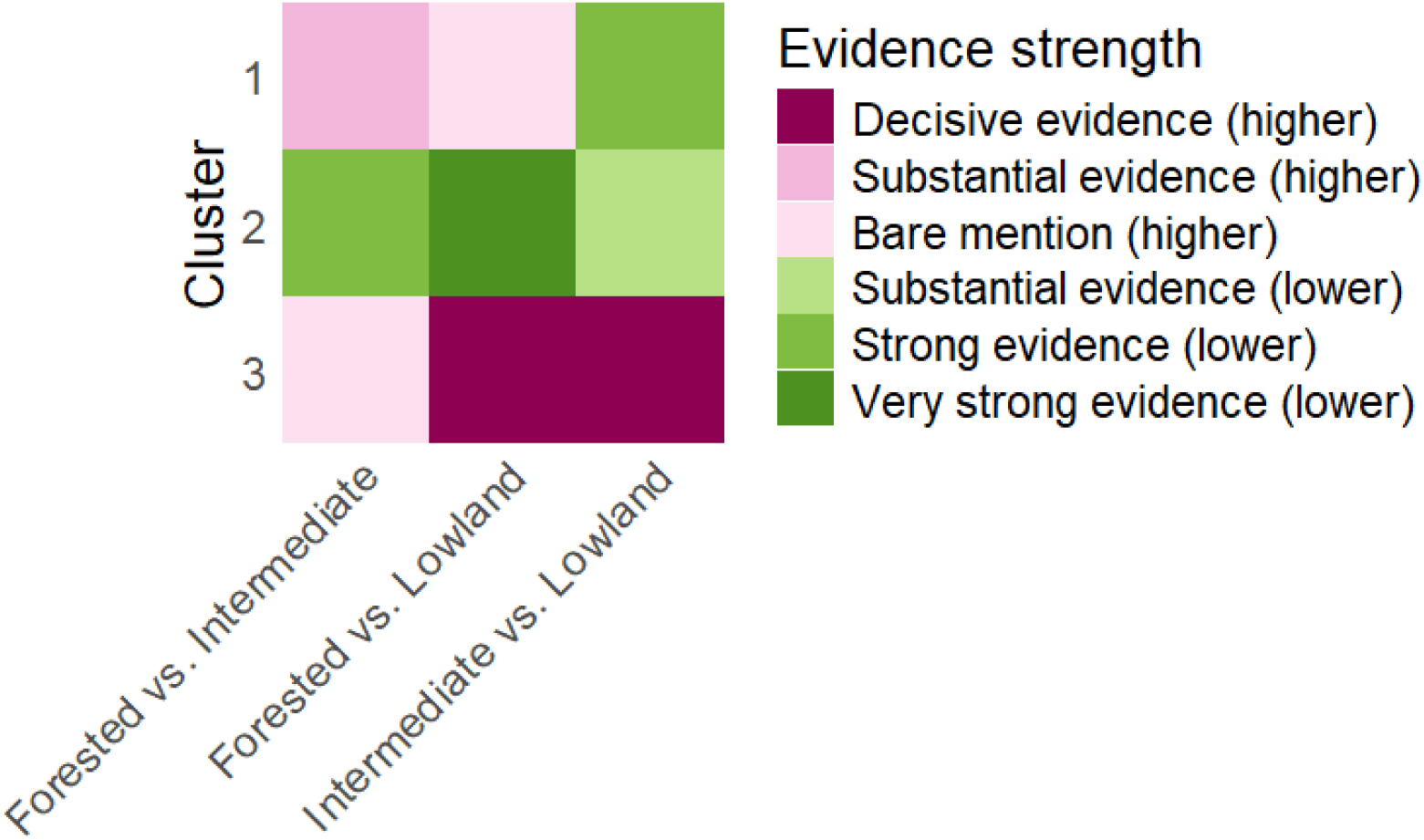
Pair-wise comparisons of straw removal rates between regions by Cluster (respondent profile) (n=92). Evidence strength according to Jeffreys (1939).

**Table 5.** Estimated proportions of respondents who did remove straw in 2020 by region and cluster membership with 95% credible intervals.

| Cluster | Forested<br>(estimated frequency<br>[CI95]) | Intermediate<br>(estimated frequency<br>[CI95]) | Lowland<br>(estimated frequency<br>[CI95]) | N<sub>total</sub><br>(per cluster) |
| --- | --- | --- | --- | --- |
| 1 | 0.50 [0.10–0.91] | 0.58 [0.24–0.90] | 0.62 [0.41–0.82] | 33 |
| 2 | – | 0.74 [0.47–0.95] | 0.39 [0.22–0.57] | 38<sup>a</sup> |
| 3 | 0.69 [0.33–0.98] | 0.76 [0.55–0.94] | 0.88 [0.70–1.00] | 19<sup>a</sup> |

<sup>a</sup> Information on one regional membership is missing.

Among respondents who did remove, net positive impacts of removing straw were, on average, expected only for farm profitability and the establishment of subsequent crops (Table 6). However, despite the negative average net values of expected impacts on all other goals, 59 respondents (63%) still decided on removal.

**Table 6.** Average partial NVEI for each goal and respondents who removed cereal and oilseed straw during 2020.

| *Goal* | *Average partial NVEI among respondents who removed the straw* |
| --- | --- |
| Farm profitability | 0.57 |
| Soil structure | -2.29 |
| Humus content effect | -2.33 |
| Establishment of the subsequent crop | 0.38 |
| Farm workflow | -0.79 |
| Tilling properties | -0.93 |
| Weed management | -0.68 |

## 4. Discussion

This study provides new insights into farmer decision-making on straw management by shifting the focus from simple adoption rates to the underlying motivations and expectations that guide behaviour. While much of the existing literature models biomass provision primarily as a question of uptake likelihood, identifying key drivers of willingness to supply biomass (e.g., Hoque et al., 2015), our approach emphasises how farmers’ expectations of outcomes (NVEI) vary across distinct respondent profiles and shape straw management decision-making. By explicitly identifying the communication needs of different farmer clusters, we provide a more nuanced understanding of how advisory efforts can be effectively tailored in practice.

Our results reveal that straw management decision-making is influenced by multiple factors, including expected effects on profitability, soil health, workflow, and crop establishment, as well as regional context. Building on agent-based frameworks, such as those presented by Pralhad et al. (2021), we explicitly quantified farmers’ expected utilities rather than relying on assumed preferences, thereby enabling us to capture the heterogeneity in farmer decision-making based on their own assessments of anticipated outcomes. Consequently, identical behaviours, removing or not removing straw, may stem from very different considerations depending on the farmer and their local context, underscoring the need for communication and policy measures that are tailored to the motivations and concerns of each farmer cluster.

It should be noted that our data were collected in 2021, before the implementation of incentives aimed at increasing carbon storage in soils and biomass under Land Use, Land Use Change, and Forestry (LULUCF) regulations (EU, 2023b), including carbon credit schemes. These policies may affect future conditions for straw removal and the relative importance of different decision-making drivers.

As in many studies on farmer decision-making and land-use preferences (e.g., Rommel et al. 2022), our survey achieved a relatively low response rate. Although this constrains the statistical generalisability of the results, previous research highlights that low participation is typical in studies addressing novel or long-term land-use changes. Nevertheless, the emergence of distinct patterns across regions and decision types indicates that the data still offer meaningful insights into farmer motivations and potential barriers.

We also recognise the potential interdependence between our auxiliary hypothesis, namely, that NVEI reliably predicts straw removal decision-making as confirmed by the BART model (Figure S1), and our main hypotheses on cluster-specific motivations and communication needs. By first establishing NVEI’s predictive relevance, we integrated this dependency into our subsequent interpretation of the main results. This approach allowed us to interpret behavioural variation not only as a matter of adoption likelihood but as a function of differentiated expectations about risks and opportunities.

The following hypotheses were used to structure the analysis and interpretation:

### 4.1 H.1 The likelihood of straw harvest removal is higher among farmers who apply manure compared to those who do not

To test H1, which posits that the likelihood of straw harvest removal is higher among farmers who apply manure compared to those who do not, we used a permutation test within a multivariate BART model. The results show that the predictor *Manure* is significantly associated with removal decision-making (p < 0.002; Figure 3). The SHAP summary plot (Figure 4) further indicates that manure application is the most influential predictor, followed by *NVEI* and *Lease share*, and confirms that farmers applying manure are more likely to remove straw than those who do not. Thus, H.1 was corroborated.

We also explored whether sludge (sewage sludge or digestate) influenced straw removal decisions. It was not a significant predictor of removal probability and was dropped from the multivariate BART model. This suggests that, unlike manure application, the use of humus-rich sludge does not strongly drive farmers’ straw management decisions. The lack of effect may reflect limited on-farm adoption, uncertainty about impacts, or knowledge gaps regarding its interaction with soil fertility. While not central to our hypotheses, these findings highlight an area where further research or advisory support could clarify the role of alternative organic amendments in sustainable straw management.

### 4.2 H.2 A higher proportion of leased land is positively associated with straw harvest removal decision-making

Blennow et al. (2025a) reported that farmers who lease most of their cultivated land are more willing to sell cereal and oilseed straw than those who farm predominantly owned land. They interpreted this as reflecting lower concern for long-term soil fertility among tenants, whose tenure is typically shorter.

In our study, a Bayesian proportion test initially suggested the opposite pattern: farmers who did not remove straw in 2020 were more likely to lease more than 50% of their cultivated land than those who did remove (Table 3). To test H.2, which posits that a higher proportion of leased land is positively associated with straw harvest removal decision-making, we employed a permutation test using the multivariate BART model, which accounted for potentially confounding factors. The results are consistent: *Lease share* is significantly associated with straw removal (p =0.002; Figure 3), but in contrast to H.2, a higher proportion of leased land is linked to a lower probability of cereal and oilseed straw removal (Figure 4). Thus, H.2 was not corroborated.

This divergence from earlier findings underscores an important distinction: a positive attitude toward straw selling does not necessarily translate into actual straw removal behaviour, as constraints may prevent farmers from acting in accordance with their stated attitudes. A comparable gap between attitudes and decisions was reported by Blennow et al. (2025c) regarding farmers’ willingness versus their actual decision-making to cultivate *Populus* spp. on agricultural land.

### 4.3 H.3 Farmers managing humus-rich soils are more likely to engage in straw harvest removal decision-making than farmers managing soils with lower humus content

To test H.3, which posits that farmers managing humus-rich soils are more likely to engage in straw harvest removal decision-making, we applied a permutation test within the multivariate BART model. The results show that *Humus content* is significantly associated with straw removal (p = 0.01; Figure 3). However, SHAP values revealed that straw removal was most likely among farmers with intermediate soil humus levels, compared with those managing soils of low or high humus content, as well as those uncertain of their soil’s humus status (Figure 4). Consequently, H.3 was not corroborated.

Nevertheless, self-reported measures such as the questionnaire responses used in this study should be interpreted with caution. Participants may provide answers that construct a consistent narrative of their decision-making, rather than reflecting independent expectations and actions. This raises uncertainty about the direction of causality: expectations might shape decisions, but decisions may equally shape expectations. Such post-decision rationalisations are well documented in behavioural research (cf. Ariely and Norton 2007) and are plausible both for farmers who had already decided to remove straw and for those who had decided against it.

### 4.4 Policy implications

Our findings have several implications for policymakers, advisory bodies, and extension services aiming to support informed decisions about straw removal and management. These insights can guide the development of communication strategies, advisory programs, and potential policy instruments, while adhering to principles of transparency and respect for farmers’ autonomy.

#### 4.4.1 Guidelines for effective communication

Based on our results, we propose the communication guidelines to prioritise informed and autonomous decision-making rather than persuasion. Efforts should address knowledge gaps, particularly regarding soil health, nutrient cycling, and farm operational outcomes. Straw removal is context-dependent and may conflict with long-term soil fertility, ecosystem services, or other agronomic priorities.

The three distinct farmer profiles identified in our study highlight specific knowledge gaps and indicate where communication can enhance farmers’ understanding of sustainable straw management. Regional variations further emphasise the need for context-specific guidance (Table 7).

**Table 7.** Communication guidelines (based on. **Figures 5–7; Tables 4–6).**

| *Profile* | *Knowledge & practices* | *Expected impacts of straw removal* | *Regional distribution* | *Communication guideline* |
| --- | --- | --- | --- | --- |
| 1 | Knows soil humus content; applies manure | Expects negative impacts on soil health and workflow; removal decision-making driven by expected positive effects on profitability, subsequent crops, and soil structure | All three regions | Emphasise safe straw removal rates; explain how manure mitigates negative impacts; highlight preservation of humus, soil structure, and long-term fertility |
| 2 | Knows soil humus content; does not apply manure | Expects negative impacts on soil health; decisions influenced by less severe expected effects on tillage and workflow | Intermediate & Lowland regions | Highlight increased risk of long-term soil degradation without manure or other supplements; stress balancing removal with practices that protect soil organic matter |
| 3 | Often unaware of soil humus content; applies manure | Expects positive impacts, primarily on subsequent crop establishment | All three regions | Focus on why it is important to know soil humus content before providing guidance on monitoring humus levels; emphasise long-term soil health alongside short-term productivity |

By identifying the communication needs of distinct farmer clusters, advisory bodies can tailor strategies that reflect not only diverse motivations and structural constraints but also local conditions. Crucially, positive attitudes toward straw removal do not automatically translate into practice; effective guidance and policy must account for structural, agronomic, local, and informational factors that jointly shape actual behaviour.

## 5. Conclusions

Farmers’ perspectives on straw harvest removal are shaped by agronomic considerations and by practices that have developed in response to local conditions, while also being influenced by broader environmental and socio-economic changes. Knowledge gaps, particularly regarding soil humus content, contribute to the diversity of perspectives.

For policy, uniform assumptions about straw availability are problematic. Strategies to mobilise biomass must account for diverse farm-level practices and knowledge, while communication should provide balanced, evidence-based information that helps farmers weigh short-term benefits against long-term impacts on soil fertility, carbon cycling, and ecosystem services. Recognising the gap between stated willingness and actual behaviour is crucial to ensure that policies and advisory programs are both realistic and effective. These findings are particularly relevant to the European Green Deal, the EU Soil Strategy for 2030, and renewable energy targets, where biomass is often treated as a homogeneous resource.

The methodology used here offers a framework for capturing the diversity of farmers’ perspectives on land management. Applying similar approaches across regions and production systems can improve understanding of decision drivers, identify knowledge gaps, and support communication and policy interventions that align biomass supply with sustainable land management.

## 6. Acknowledgements

We gratefully acknowledge all interviewees and questionnaire participants in Scania County for their invaluable input. Our thanks also go to the Swedish Farmers Association for supporting the participation of potential respondents, and to the Swedish Board of Agriculture for granting access to relevant data.

## Data availability

The data can be accessed from the Swedish National Data Repository () by anyone with a legitimate interest in the data, as long as the transfer of data complies with the Swedish and European regulations on data protection.

