## Supplemental Information for "Beyond adoption rates: Farmer motivations and communication needs in straw management decision-making"

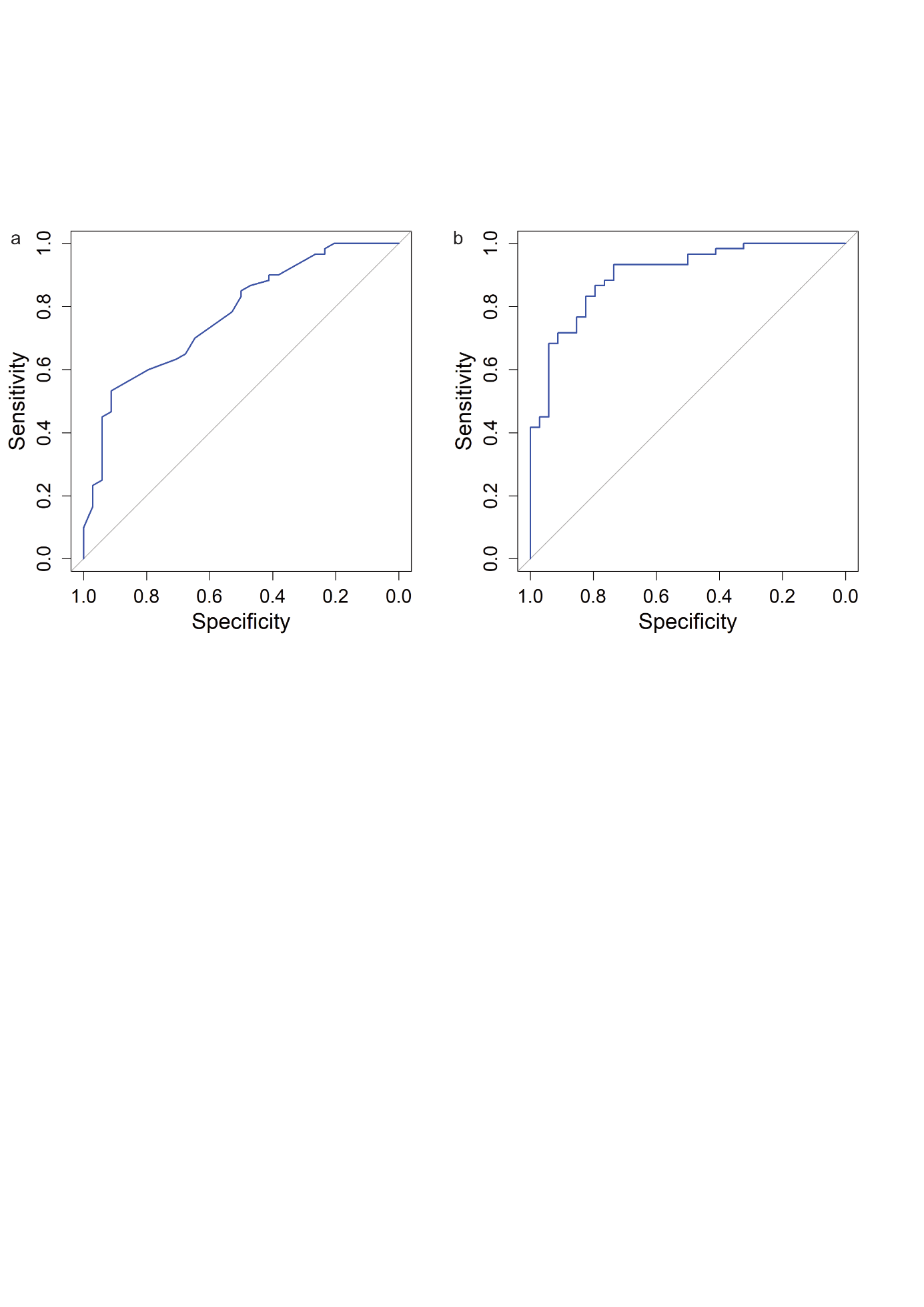

Figure S1. Receiver Operating Characteristic (ROC) curve for the univariate (a) and multivariate (b) BART model, with an Area Under the Curve (AUC) of 0.77 and 0.90, respectively.

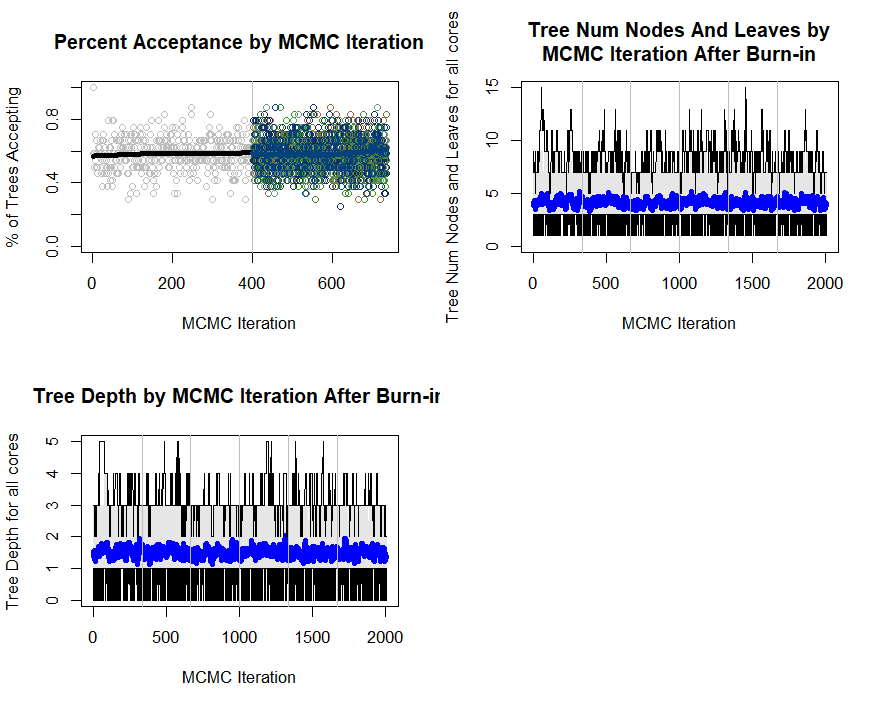

Figure S2. Convergance of multivatiate BART model.
